# The molecular basis of phenotypic plasticity evolves in response to environmental predictability

**DOI:** 10.1101/2022.10.31.514467

**Authors:** Christelle Leung, Daphné Grulois, Leandro Quadrana, Luis-Miguel Chevin

## Abstract

Phenotypic plasticity, the response of a given genotype to its environment of development, is a ubiquitous feature of life, enabling organisms to cope with variation in their environment. Theoretical studies predict that, under stationary environmental variation, the level of plasticity should evolve to match the predictability of selection at the timing of development. However, we still lack critical empirical evidence on the extent to which selection on phenotypic plasticity cascades down from higher phenotypic levels to their underlying molecular basis. Here, we used experimental evolution under controlled environmental fluctuations, to test whether the evolution of phenotypic plasticity in responses to environmental predictability (*ρ*^*2*^) occurred across biological levels, going from DNA methylation to gene expression to cell morphology. Transcriptomic results indicate clear effects of salinity and *ρ*^*2*^×salinity interaction on gene expression, thus identifying sets of genes involved in plasticity and its evolution. These transcriptomic effects were independent of DNA methylation changes in *cis*. However we did find *ρ*^*2*^*-*specific responses of DNA methylation to salinity change, albeit weaker than for gene expression. Overall, we found consistent evolution of reduced plasticity in less predictable environments for DNA methylation, gene expression, and cell morphology. Our results provide the first clear empirical signature of plasticity evolution at multiple levels in response to environmental predictability, and highlight the importance of experimental evolution to address predictions from evolutionary theory, as well as investigate the molecular basis of plasticity evolution.

## 1. Introduction

Phenotypic plasticity, the ability of a given genotype to produce different phenotypes depending on environmental conditions, is an important mechanism enabling organisms to cope with variation in their environment. Understanding what drives the evolution of plasticity, from its selective causes to its underlying mechanisms, is thus important not only for basic research, but also for on our ability to predict the fate of populations, especially under global change [1, 2]. Despite this interest, we still lack critical empirical information on the extent to which selection on phenotypic plasticity propagates across hierarchical levels of the organisms, from “higher phenotypes” that are directly exposed to selection, to their underlying molecular basis.

Theoretical and empirical studies have demonstrated that the adaptiveness of phenotypic plasticity arises from its interplay with environmental variation in selection, when the latter is partly predictable [3-7]. A high degree of plasticity is expected to be favored when the environmental cue causing induction of a given phenotype is a reliable predictor of selection acting on this phenotype. Conversely, under unpredictable environmental changes (or unreliable cues), plasticity leads to phenotypes that are often mismatched with their optimum, such that lower degrees of plasticity are selected [3-7]. In the limit of completely unpredictable environments, bet-hedging strategies that do not rely on environmental cues may even be selected [8, 9].

In terms of mechanisms and pathways, the increased availability of -*omics* methods has greatly facilitated our understanding of the relationship between molecular changes and changes in phenotypic expression. In particular, many studies have shown that phenotypic plasticity often involves at its core variation in gene expression with the environment [10-14]. This suggests that the molecular basis of phenotypic plasticity should also encompass mechanisms that regulate gene expression, as clearly demonstrated in a number of cases [12-14]. In particular epigenetic processes, that is, modifications of chromatin beyond DNA sequence, transmitted through mitosis (and possibly meiosis) and potentially influencing gene expression, were shown to be important molecular mechanisms for phenotypic plasticity [15-18]. As both gene expression and epigenetic marks (such as DNA methylations) are known to be under genetic control [19-21], selection on phenotypic plasticity for higher traits is thus expected to cascade down to cause evolution of plasticity for its underling molecular mechanisms [e.g. 22], but this process has seldom been investigated experimentally. More specifically, the extent to which classical theoretical predictions about evolution of plasticity in response to environmental predictability hold from molecular phenotypes to more integrated ones remains unexplored.

Recently, we have showed that environmental variation in CpG methylation and gene expression in the halotolerant microalga *Dunaliella salina* contributes to phenotypic plasticity in this species [23]. In addition, we confirmed the contribution of the genotype to salinity responses at the levels of DNA methylation, gene expression, and growth rate, highlighting the evolutionary potential of phenotypic plasticity at multiple levels [23]. Importantly, long-term experimental evolution with this species has demonstrated that reduced plasticity in cell shape and content evolved in populations confronted to less predictable environments [5]. Here, we used experimental evolution under controlled environmental fluctuations to assess whether the evolution of plasticity in responses to environmental predictability permeates across biological levels, from DNA methylation to gene expression to higher phenotypes.

## 2. Results

We analyzed nine populations of the halotolerant microalga *Dunaliella salina* (strain CCAP 19/15) that have evolved under regimes of randomly fluctuating environments, with controlled and variable predictability [5, 24]. During experimental evolution, lines derived from a single ancestral population were exposed to randomly fluctuating salinity, with changes every three or four generations, for a total of *c*. 500 generations. Salinity had a normal distribution over time, with the same mean (2.4M NaCl) and standard deviation (1M NaCl) across treatments, but variable autocorrelation, and hence variable predictability [5, 24]. Previous morphological analysis of 32 of these lines revealed that reduced morphological plasticity has evolved in lines that experienced less predictable environments [5]. To further characterized the molecular basis of this evolution, we set out to determine whether the plasticity of DNA methylation and gene expression levels for a subset of these lines evolved under three different predictability treatments (three lines per treatment): low (*ρ*^*2*^ = 0), intermediate (*ρ*^*2*^ = 0.25) and high (*ρ*^*2*^ = 0.81) predictability, where *ρ* is the stationary (long-term) temporal autocorrelation of salinity time series. All populations were subsequently subjected to a ten-day acclimation step at intermediate salinity ([NaCl] = 2.4 M), to ensure they had similar physiological states and population densities before the phenotypic and molecular assays. They were then placed for 24 h at two salinities near the extremes of their historical range ([NaCl] = 0.8 M and 4.0 M), to assess their degree of plasticity in DNA methylation, gene expression and individual cell morphology (Fig. 1).

**Figure 1.**
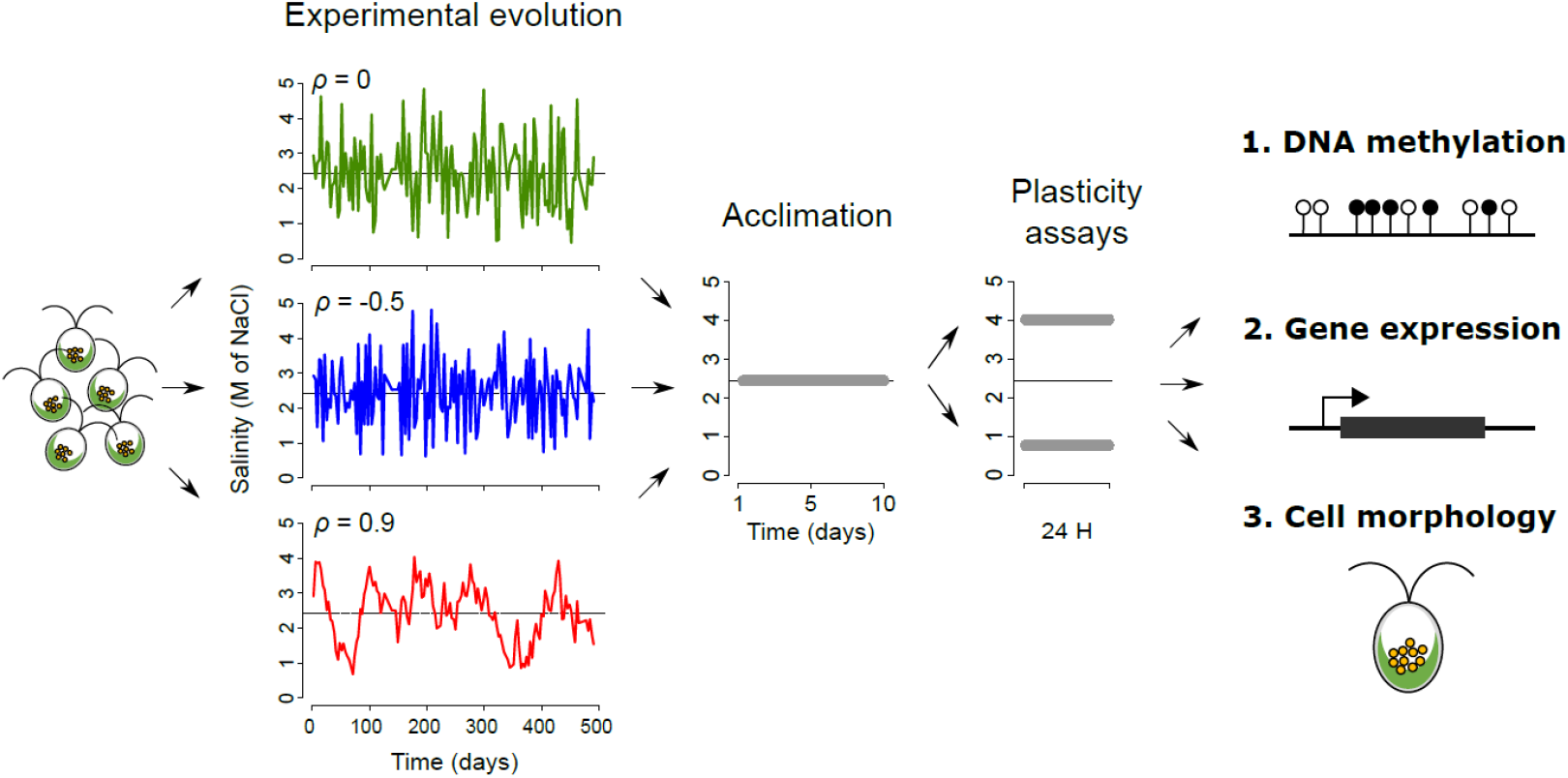
Experimental design. The experiment included three steps: (i) long-term experimental evolution (left); (ii) acclimation at a constant, intermediate salinity (middle); and (iii) plasticity assays at high versus low salinity (right). Each colored time series on the left represents an actual realization of salinity fluctuations for one of the populations used in this study, with the color denoting the treatment of stationary (that is, expected long-term) temporal autocorrelation (*ρ*). At the end of the experiment, cells were harvested for DNA methylation, gene expression and cell morphology analyses.

### 2.1. DNA methylation and gene expression plasticity evolved in response to environmental predictability

DNA methylation can contribute to morphological differentiation, by influencing gene regulation [18, 25-28]. To assess whether experimental evolution of *Dunaliella salina* may lead to epigenetic differentiation, we performed whole-genome bisulfite sequencing (WGBS) for all samples, yielding a total of 1.23 × 10^9^ 150 bp paired-end raw reads, and an estimated average depth of coverage of 43.76 × (*s.d*. 3.14 ×) per sample (Table S1). After data filtering, we carried out our methylation analyses on an average of 7.56 × 10^7^ (*s.d*. 8.28 × 10^6^) cytosines per samples at the CpG context (Table S1), where methylations are most prevalent in this species [23]. Redundancy analyses based on overall CpG methylation revealed a significant effect of evolutionary treatments *ρ*^*2*^ (*R*^*2*^_*adj*_ = 4.66%; *P* = 0.005) on DNA methylation (Fig. 2A). However, we did not detect a significant marginal effect of salinity (*R*^*2*^_*adj*_ = 0.79%; *P* = 0.356) or *ρ*^*2*^ × salinity interaction (*R*^*2*^_*adj*_ = 0.30%; *P* = 0.562) on the overall DNA methylation pattern.

**Fig. 2.**
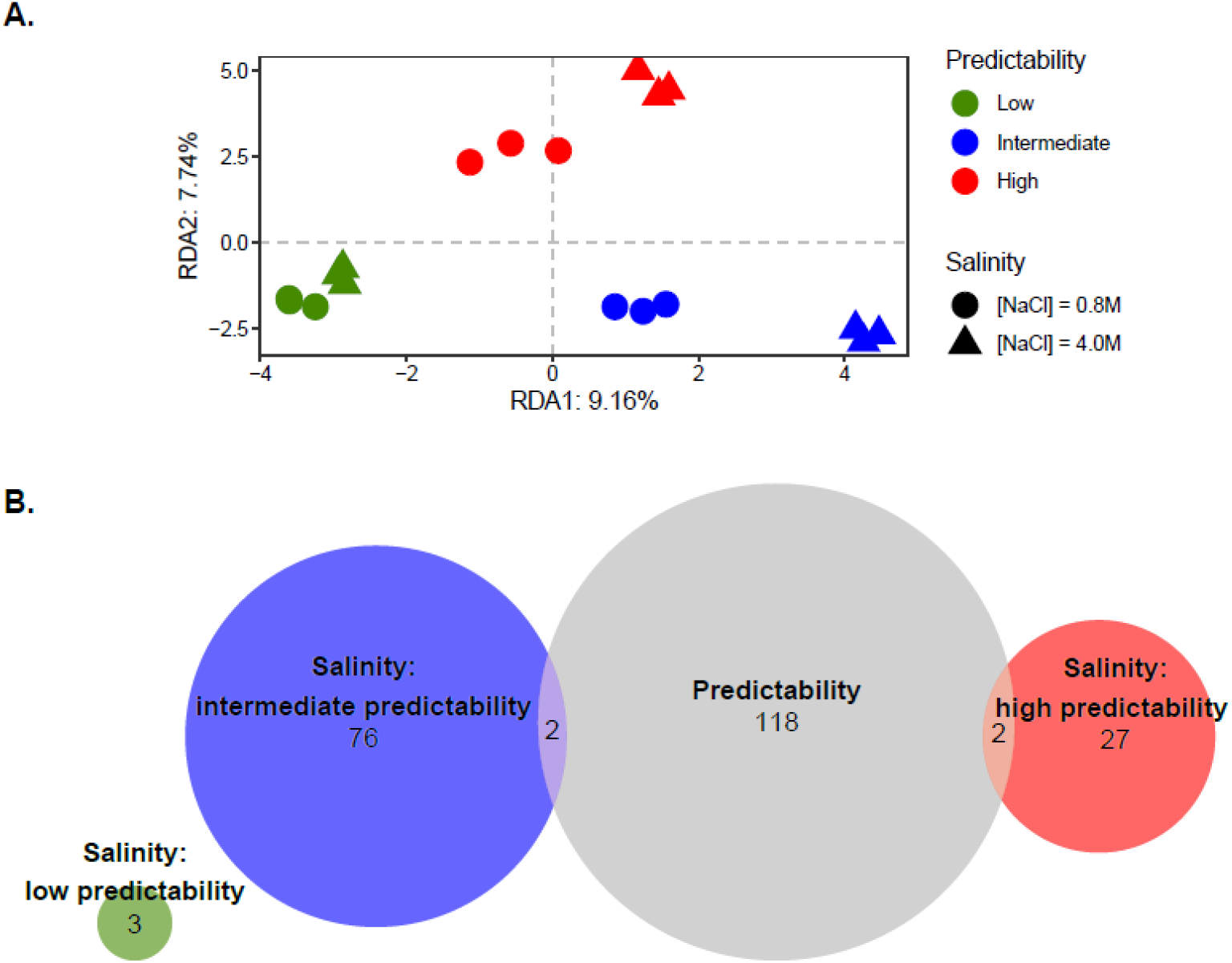
Evolution and plasticity of DNA methylation. **A**. Variation of DNA methylation patterns across evolved populations. Redundancy analysis (RDA) plot performed on DNA methylation patterns according to evolution conditions (colours) and salinity during plasticity assay (shapes). **B**. Distribution of differentially methylated regions (DMRs). Venn diagram describing the number of DMRs (*q*-value < 0.05 and |diff-Methylation| > 20%) across evolutionary conditions (Predictability, light grey), and between salinities within each of the three evolutionary conditions: low (green), intermediate (blue) and high (red) predictability of environmental changes.

Lack of significant differentiation at the whole epigenome level does not preclude more localized epigenetic differences, so we also investigated regional changes in DNA methylation at a finer scale, by considering non-overlapping 100 bp windows, hereafter denoted Differentially Methylated Regions (DMRs). We detected 227 DMRs among the evolutionary treatments, as summarized by their environmental predictability *ρ*^*2*^ (Fig. 2B). We also detected *ρ*^*2*^-specific DMRs between salinities (*q*-value < 0.05 and |diff-Methylation| > 20%) within each evolutionary treatment, among the 14,357 total 100bp regions (Fig. 2B). Interestingly, populations that evolved under less predictable environmental fluctuations displayed the least number of DMRs between salinities (n = 3), as compared to populations from intermediate (n = 78) or high (n = 29) environmental predictability (Fig. 1B), indicative of reduced epigenetic plasticity. We then assessed whether changes in DNA methylation patterns in response to a given environmental challenge involved similar genomic regions in the different evolved lines. Comparison of the list of DMRs between salinities revealed no overlap across evolutionary conditions (Fig. 2B), suggesting that evolution of plastic epigenetic responses involved modifications of methylations in distinct genomic regions in different treatments.

We next investigated variation in gene expression, by analyzing 32718 transcripts through RNA-sequencing. We obtained 5.62 × 10^8^ 150 bp paired-end raw reads in total (Table S2). As for DNA methylation, we detected a significant effect of the evolutionary treatments *ρ*^*2*^ (*R*^*2*^_*adj*_ = 9.79%; *P* < 0.001) on gene expression at the whole-transcriptome level. However, the assay salinity now explained the greatest part of variation in gene expression (*R*^*2*^_*adj*_ = 32.91%; *P* < 0.001), indicating pervasive and highly significant plasticity. The *ρ*^*2*^ × salinity interaction was also significant (*R*^*2*^_*adj*_ = 4.30%; *P* = 0.029), (Fig. 3A) indicating evolution of transcriptional plasticity. At a more local level, analyses of differentially expressed (DE) transcripts yielded similar results: the numbers of transcripts that were significantly differentially expressed (Likelihood ratio test, FDR < 0.05 and |Log_2_FC| > 1) was highest for contrasts between salinities (n = 4,283), followed by evolutionary treatments *ρ*^*2*^ (n = 1,315), and finally *ρ*^*2*^ × salinity interaction (n = 837). As for DMRs, populations that evolved in less predictable environments displayed fewer DE transcripts than populations that evolved in highly predictable environments (Fig. 3B, n = 2638, 2844 and 4086 for *ρ*^*2*^ = low, intermediate and high, respectively). However, in contrast to what we found for DNA methylation, we observed a great overlap among DE transcripts identified between salinities for the different evolved lines (Fig. 3B), indicating that the plastic response to salinity largely involved transcriptional regulation of a common pool of genes, regardless of their evolutionary trajectory. Nonetheless, we still detected some *ρ*^*2*^-specific DE transcripts between salinities (Fig. 3B).

**Fig. 3.**
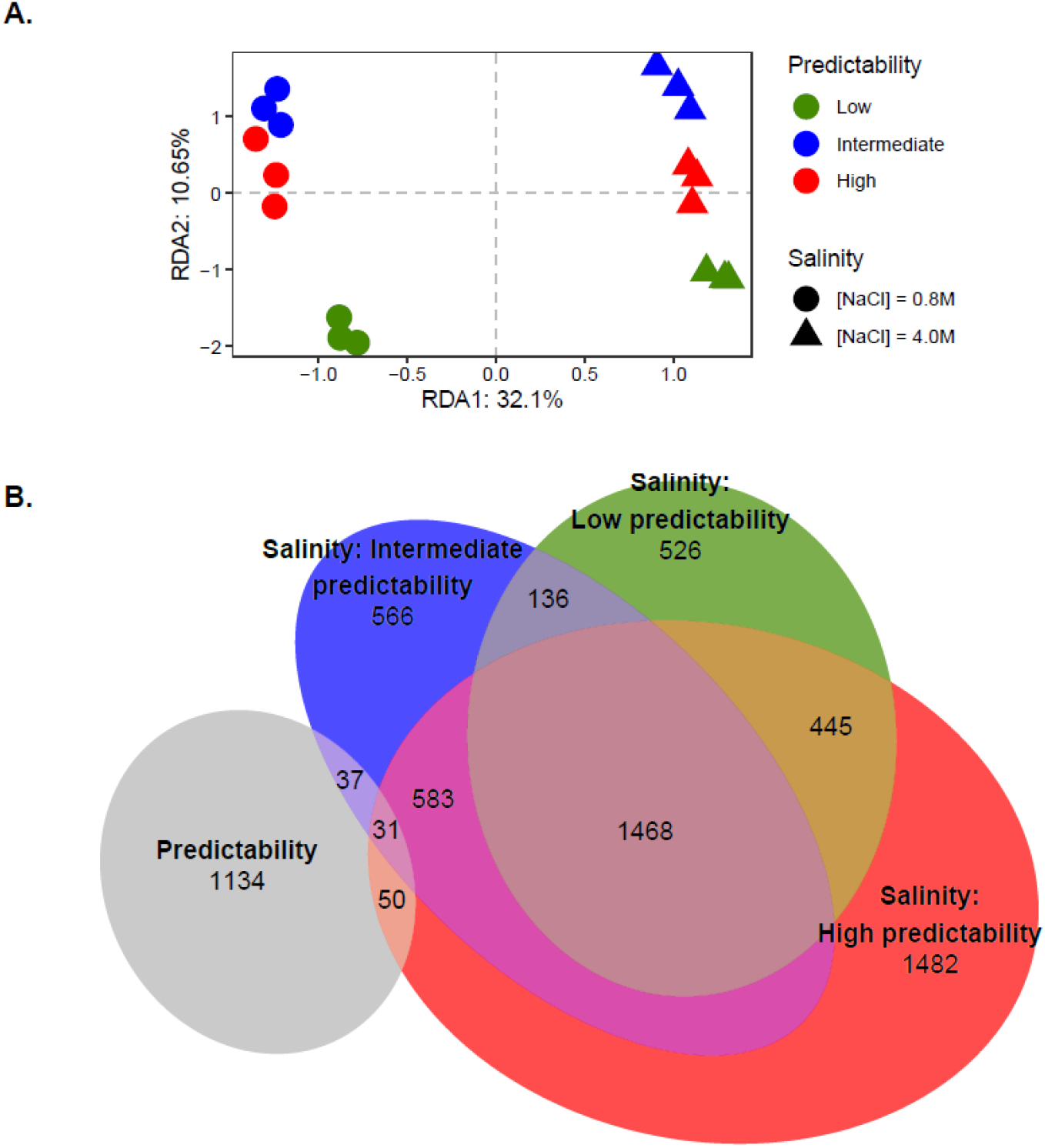
Evolution and plasticity of gene expression. **A**. Variation of gene expression levels across evolved populations. RDA plot performed on gene expression levels according to evolution conditions (colours) and salinity during plasticity assay (shapes). **B**. Distribution of differentially expressed transcripts. Venn diagram describing the numbers of differentially expressed transcripts across evolutionary conditions (Predictability, light grey) identified by performing likelihood-ratio test (LRT, FDR < 0.05) as implemented in *DESeq2*, and between salinities (Wald test, FDR < 0.05 after BH adjustment and |log_2_FC| > 1) for three evolutionary conditions: low (green), intermediate (blue) and high (red) predictability of environmental changes.

To confirm that differences in gene expression (and to a lesser extent DNA methylation) among salinities were due to plasticity, rather than resulting from putative strong selection taking place in the polymorphic populations during the short duration of the assay, we also sequenced isogenic populations (i.e. founded from a single, presumably haploid cell from the evolved populations) subjected to different salinities. Despite the genetic homogeneity of these population, we confirmed the salinity effect on both DNA methylation and gene expression (Fig. S1).

We next performed Gene Ontology (GO) enrichment analysis, to assess gene functions involved in *D. salina* response to salinity changes and its evolution, using the functional genome annotation constructed in Leung, et al. [23]. Transcript functional annotation analysis revealed that DE transcripts between salinities mostly involved genes associated to cellular components and molecular functions involving the chloroplast and protein transport (Fig. 4). While no GO term enrichment was detected for DE transcripts among evolutionary treatments (for FDR < 0.1), we were able to detect that *ρ*^*2*^ × Salinity interaction effects on gene expression essentially involved gene functions associated different metabolic processes within the GO category ‘biological process’ (Fig. 4).

**Fig. 4.**
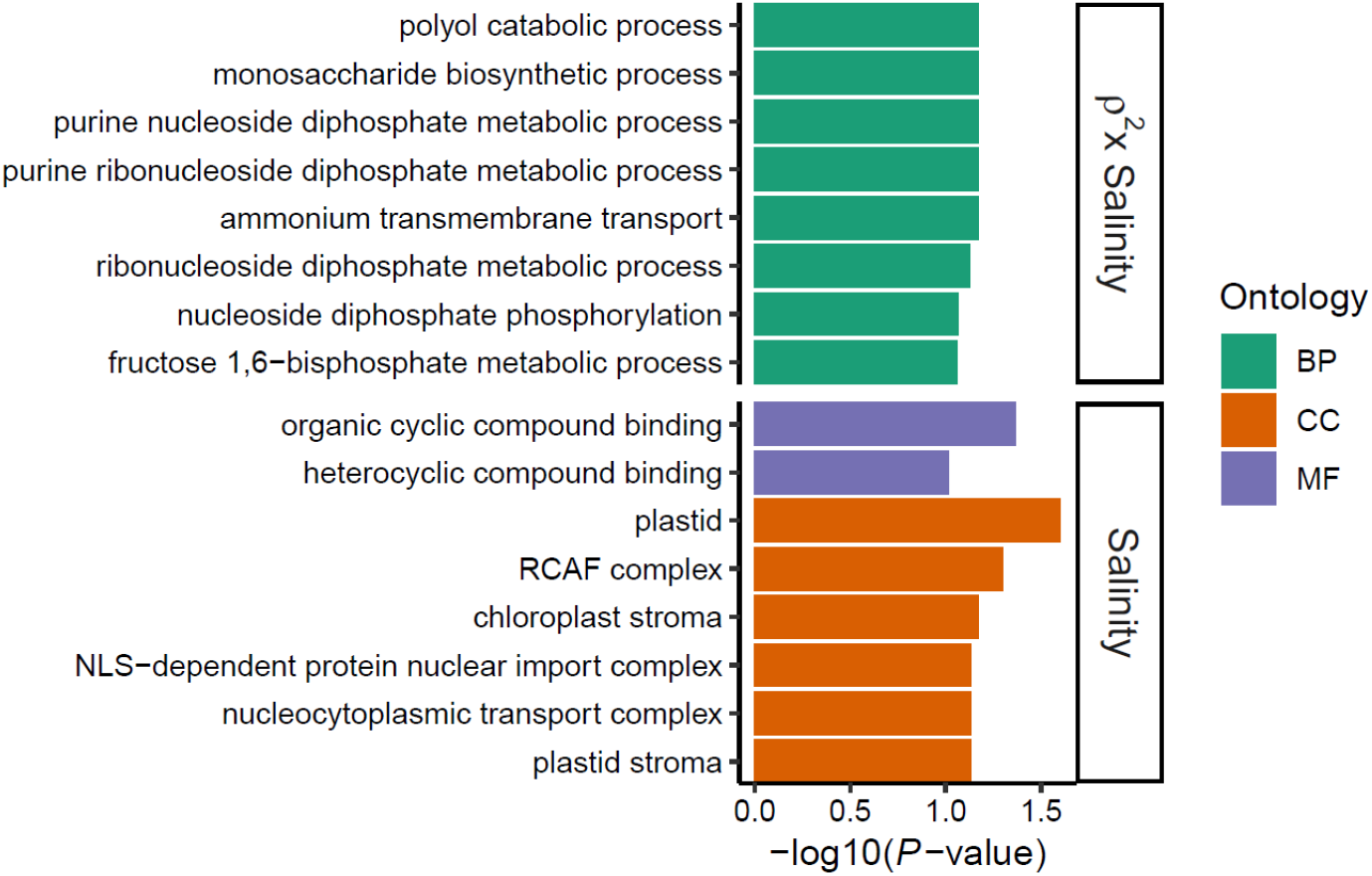
Gene functions involved in salinity-induced plasticity and its evolution. Enriched GO terms of identified DE transcripts between salinities (Salinity) and *ρ*^*2*^-specific transcriptional response (*ρ*^*2*^ × Salinity). GO categories included molecular function (MF, purple), cellular component (CC, orange) and biological process (BP, green), and were sorted by decreasing order of evidence within each category based on GO enrichment test P-value (for FDR ≤ 0.1). No GO term enrichment with FDR ≤ 0.1 was detected for DE transcripts among evolutionary treatments (*ρ*^*2*^).

### 2.2. DNA methylation in cis were not associated with gene expression

We assessed whether DNA methylation could influence the expression of neighbouring genes in *cis*. For each methylated cytosine, we searched for the nearest transcription start site (TSS). For this analysis, all cytosines associated to the same transcript were merged into a common gene-associated methylation region, following the same method used in Leung, et al. [23]. Among the 3,064 methylated regions associated to a transcript, we detected 18 regions that were significantly differentially methylated among evolutionary treatments (including all comparisons of *ρ*^*2*^ pairs), and 21 regions among salinities (for each evolutionary treatments *ρ*^*2*^). However, only four of these 39 transcript-associated regions were associated to significant DE transcripts for the same comparison, and only one involved a comparison between salinities. This suggested that, should DNA methylation have an effect on gene expression plasticity, this effect must be acting on regulation in *trans*.

### 2.3. The degree of plasticity evolved consistently across levels

We detected significant effect of salinity (*R*^*2*^_*adj*_ = 2.51%; *P* < 0.001), evolutionary treatment *ρ*^*2*^ (*R*^*2*^_*adj*_ = 4.13%; *P* < 0.001), and their interaction (*R*^*2*^_*adj*_ = 1.06%; *P* < 0.001) on cell morphology, thus providing independent replication of the results from Leung, et al. [5]. To investigate the consistency in the direction of experimental evolution of plasticity across levels, we then quantified the overall degree of plasticity for each of them, measured as the Euclidean distance between low and high salinities. When regressing the magnitude of plasticity against the environmental predictability of the experimental evolution treatment, we consistently found a positive relationship, with weak to moderate evidence [sensu 29] for DNA methylation (*P* = 0.057, Fig. 5A), but strong evidence for gene expression (*P* = 0.018, Fig. 5B) and cell morphology (*P* = 0.004, Fig. 5C). Hence, populations that have experienced less predictable environments during experimental evolution have evolved reduced plasticity, not only for cell morphology (as shown by Leung, et al. [5]), but also for DNA methylation and gene expression, two important molecular mechanisms potentially underlying phenotypic plasticity.

**Fig. 5.**
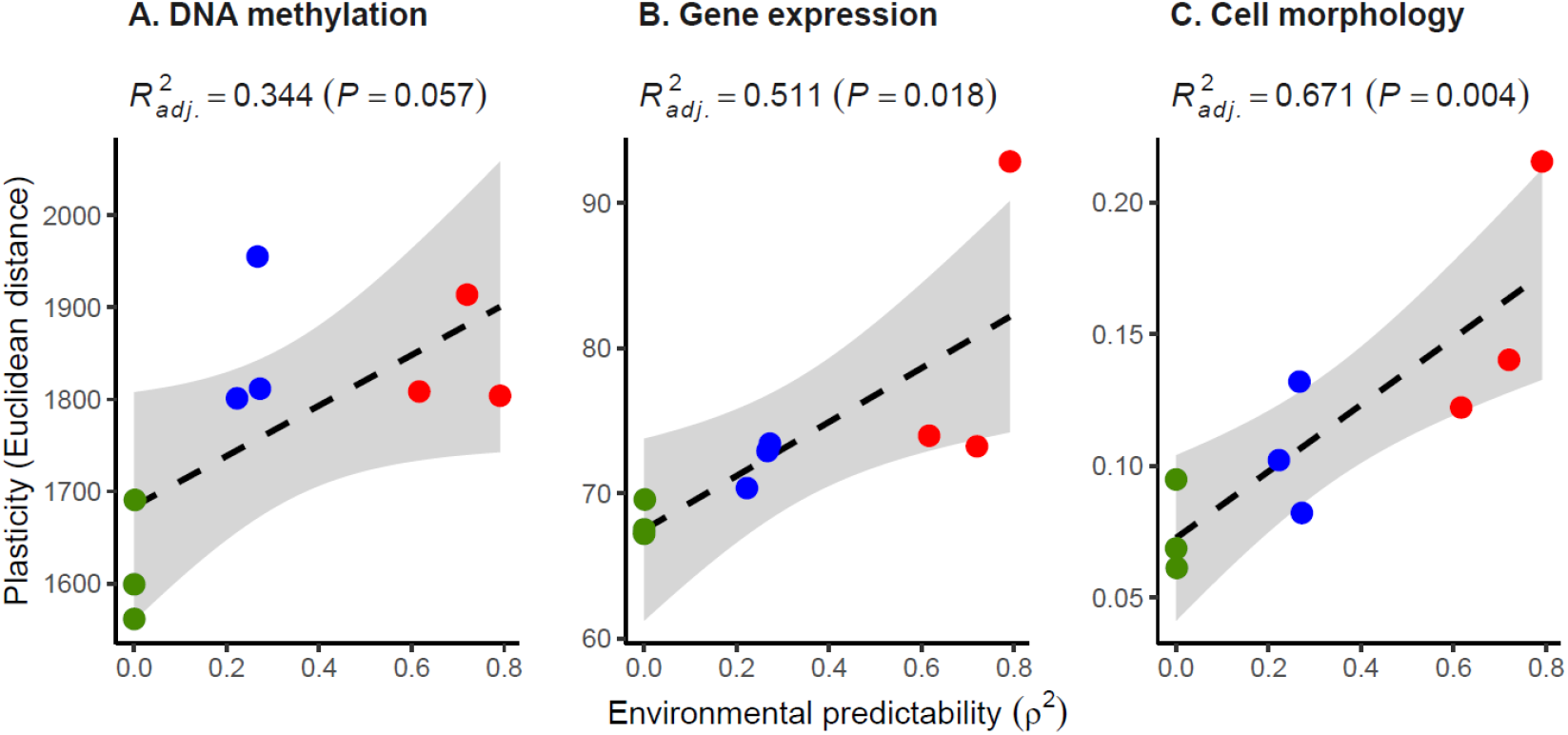
Multi-level evolution of plasticity in response to environmental predictability. The degree of plasticity of populations, as measured by the Euclidian distance between low and high salinities for (A) DNA methylation patterns at CpG context, (B) gene expression levels, and (C) cell morphology, is plotted against the predictability *ρ*^*2*^ of the environmental fluctuations that these populations have experienced during experimental evolution. The dashed line is the regression slope, and the grey area represents the 95% confidence interval of the linear regression.

## 3. Discussion

We aimed to understand whether the evolution of plasticity in response to environmental predictability is a phenomenon that can be observed at different hierarchical levels. We have therefore exposed experimental populations of the microalga *Dunaliella salina* to randomly fluctuating salinity with controlled predictability, and then assessed how their plasticity evolved at three levels: DNA methylation, gene expression, and cell morphology. Our results highlighted important aspects of the molecular mechanisms of plasticity and its evolution.

### 3.1. Gene expression plasticity and its regulation

A wealth of studies have investigated the molecular underpinnings of phenotypic plasticity, and most of them have identified gene expression as a key mechanism of phenotypic changes [10-14]. A crucial step in gene expression is transcription to mRNA, which explains a large fraction of variation in protein abundance [30], and therefore in higher phenotype. Here, we found that salinity-induced transcriptional plasticity of *D. salina* involved largely overlapping Differentially Expressed transcripts among different populations, indicating that this species holds specific genes to cope with salinity changes in its environments. In addition, we also detected significant *ρ*^*2*^ × *Salinity* on gene expression, indicative of the evolution of transcriptional plasticity with respect to environmental predictability. While only ca. 30% of the total transcripts were successfully assigned to at least one Gene Ontology term [23], we were able to assess some gene functions involved in *D. salina* response to salinity. On the one hand, common salinity responses mostly involved genes associated to chloroplast and membrane-associated functions, like protein transport. On the other hand, *ρ*^*2*^-specific salinity responses entailed biological processes necessary for the production of energy for various functions of the organisms [31-33]. From a functional standpoint, these results suggest that *D. salina* preserves its complex molecular machineries to cope consistently with osmotic stress, while evolutionary differentiation in salinity-induced plasticity may involve potential energy allocation trade-offs among different functions.

Regulation of gene expression involves different mechanisms, including epigenetic processes [34, 35]. Here, we investigated one such epigenetic processes, namely DNA methylation. We detected Differently Methylated Regions (DMRs) between salinities for every evolutionary treatments, confirming that DNA methylation is a process that could be involved in phenotypic plasticity in this species [23]. However, we failed to detect a correlation between DMRs and the expression of proximal genes, suggesting that regulation of gene expression plasticity for this species may be driven by *trans*, instead of *cis*, regulatory changes [36, 37]. Alternatively, mechanisms of gene expression regulation could be highly stress- and organism-dependent [38]. For *D. salina*, the time lapse (24h) between osmotic changes and DNA methylation measurement in our study could be too short to enable detection of significant correlation between DNA methylation and gene expression [39]. The first responses to osmotic shock in *D. salina* may implicate homeostatic regulation, relying on other mechanisms than DNA methylation [40].

Exploring the genetic basis of phenotypic plasticity and its evolution requires the identification of genes whose expression varies across environments, as well as loci contributing to this variable degree of plasticity among genotypes. Interestingly, the *ρ*^*2*^-specific DMRs for the three categories of evolutionary treatments did not display any overlap, suggesting that different regions in the genome are involved in the control of salinity-induced plasticity in this species. This could be the result of epistatic interactions, where different loci could contribute to fine-tune the phenotype towards its optimum [41-43].

### 3.2. Multi-level evolution of plasticity

To what extent do we expect evolution to take the same course at different biological levels, along the hierarchy of traits that relates the genotype to higher phenotypes and fitness? This question has received increasing attention in the evolutionary literature, where it was mostly expressed with respect to parallel/convergent evolution in response to a constant environmental challenge [44, 45]. Theoretical and empirical work has stablished that the amount of parallelism at each biological level depends on the degree of redundancy in the mapping from one level to the next [46-48]. This in turn depends on the extent to which each item at a given level is connected to one or many items at the level above; for instance, how many traits each mutation modifies (pleiotropy) [49-51]. Because mappings along hierarchies of traits are often redundant, we do not necessarily expect to find the same degree of parallelism at the level of genes as we find at the level of functional categories, organs, or highly integrated traits [44, 45]. And reciprocally, the degree of parallelism across levels conveys information about the redundancies in mappings along hierarchies of traits.

Here, we applied a similar approach to a classic topic from theoretical evolutionary ecology: the evolution of phenotypic plasticity in response to environmental predictability [3-7]. Just like for evolutionary parallelism, we do not necessarily expect the evolution of plasticity in response to environmental predictability to be consistent across levels. Indeed, it was recently highlighted that the comparison of phenotypic plasticity (and its evolution) across levels can bring precious information about constraints in the genotype-(environment)-phenotype, including - but not restricted to - the above-mentioned redundancies. For instance, this can yield insights on the extent to which mutations and environments have similar influences on phenotypic variation, and which one is likely to drive evolution of the other [52, 53]. Here, we not only found substantial plasticity at the levels of DNA methylation, gene expression, and cell morphology, but also that plasticity evolved in a consistent direction - and consistent with theoretical predictions - in response to environmental predictability across these levels. This response was however less pronounced for DNA methylation, which only displayed marginally significant relationship between total plasticity and predictability (Fig. 5A), but with substantial variation in the number of salinity DMRs among evolutionary treatments (Fig. 2B).

Such consistent evolution of plasticity across levels may indicate that many, unmeasured, “higher traits” exhibit similar evolutionary responses to environmental predictability as those we have measured, such that their transcriptomic (and to some extent epigenetic) basis is less subject to redundancies. Alternatively, selection may operate to some extent independently at each hierarchical level. For instance, expression plasticity of a given gene may partly influences fitness by itself, rather than only via its effect on the plasticity of higher phenotypes. Investigating these questions would require mechanistic studies beyond the scope of this work. Nevertheless, our finding that plasticity can experimentally evolve at multiple levels in response to environmental predictability, and in a consistent direction, brings much needed empirical insights into the mechanisms underlying plasticity evolution, and highlights the usefulness of experimental evolution for investigating ambitious questions at the forefront of evolutionary biology.

## 4. Material and Method

### 4.1. Experimental evolution conditions and plasticity assays

To investigate the evolution of plasticity in responses to environmental predictability at the molecular level, we analysed different populations of the *Dunaliella salina* strain CCAP 19/15 that evolved in a fluctuating environment for *c*. 500 generations, from the experiment described in Rescan, et al. [24] and Leung, et al. [5]. Briefly, different populations starting from the same ancestor were exposed to randomly fluctuating salinity, with changes every three or four generations (i.e. twice a week, assuming one generation per day [54]). Each of these populations were subjected to independent random time series of salinity changes, over a continuous range. The time series were characterized by the same stationary mean (*μ* = 2.4 M [NaCl]) and variance (*σ* = 1), but differed in how salinity at a given time depends on the salinity prior the transfer (i.e. temporal autocorrelation *ρ* of salinity). The predictability of salinity changes for a given time series was assessed by *ρ*^*2*^, the proportion of temporal variance in salinity explained by the previous salinity, prior to the latest transfer (see Rescan, et al. [24] and Leung, et al. [5] for detailed protocol). We specifically analysed nine populations from three different target autocorrelation treatments (*ρ* = 0, -0.5 and 0.9; three populations per autocorrelation treatments). These autocorrelation treatments correspond to low (*ρ*^*2*^ = 0), intermediate (*ρ*^*2*^ = 0.25) and high (*ρ*^*2*^ = 0.81) predictability of environmental changes. While similar level of morphological plasticity was found between populations that evolved in intermediate predictability changes, but with different autocorrelation treatments (*ρ* = -0.5 and *ρ* = 0.5, Leung, et al. [5]), here we chose the negative autocorrelation include a larger range of *ρ*, thus enabling to better disentangle the effect of *ρ* and *ρ*^*2*^, if any.

Plasticity assays were performed following the protocol descried in Leung, et al. [5]. We first acclimatized all evolved lines during 10 days in the same environmental conditions ([NaCl] = 2.4 M) to ensure that all cells were in similar physiological states and at similar population densities at the beginning of the phenotypic assay. Cells grew in suspension flasks containing artificial seawater with additional NaCl to reach the required salinity, complemented with 2% Guillard’s F/2 marine water enrichment solution (Sigma; G0154–500 ML), and incubated at a constant temperature 24°C with a 12:12 h light/dark cycle with a 200 μmol.m^−2^.s^−1^ light intensity. Target salinity was achieved by mixing the required volumes of hypo-([NaCl] = 0 M) and hyper-([NaCl] = 4.8 M) saline media, accounting for the salinity of the inoculate. At the end of the acclimation step, we transferred *c*. 1 × 10^5^ cells.mL^−1^ of each populations to low ([NaCl] = 0.8 M) and high ([NaCl] = 0.8 M) salinities, for a total volume of 250 mL. After 24h following the salinity changes, we harvested the cells by centrifugation at 5,000 rpm for 15 minutes at room temperature, and cell pellets were stored at - 80°C until acid nucleic extraction.

We also measured intrinsic structural parameters of cells by passing a subsample of 150 μL of each populations through a Guava® EasyCyte™ HT flow cytometer (Luminex Corporation, TX, USA), also following the protocol described in Leung, et al. [5]. We specifically assessed the environment-specific cell morphology using the Forward Scatter (FSC) and Side Scatter (SSC) as proxies for cells size and complexity (cytoplasmic contents) [55], respectively, and fluorescence emission at 695/50 nm band pass filter (Red-B) values for chlorophyll production [56]. Cells morphology matrix consisted of values for these three parameters (FSC, SSC, and Red-B) for 150 randomly sampled cells identified as alive *D. salina* for each populations.

### 4.2. Sample preparation, sequencing, and bioinformatic preprocessing

To investigate the molecular mechanisms involved in osmotic stress responses, we performed whole-transcriptome shotgun sequencing (RNA-seq) for the comparison of gene expression levels, and whole-genome bisulphite sequencing (WGB-seq) for the comparison of DNA methylation variation among the nine evolved lines and two environmental conditions (hypo- and hyper-osmotic). The different lines started from potentially genetically diverse population, and could thus be polymorphic at the end of experimental evolution. To assess whether genetic variation within population affects the observed plasticity levels for gene expression or intra-population DNA methylation variation, we also founded three isogenic populations (one per autocorrelation treatment) started from a single cell using cells-sorting flow cytometry (BD FACSAria™ IIu; Biosciences-US). As *Dunaliella salina* is haploid, we expected all derived cells of a given population to be genetically identical. Each of these isogenic populations were also subjected to the plasticity assay as described above.

Total RNA extraction and purification of 24 samples ((9 lines + 3 isogenic populations) × 2 salinities) was carried out using Nucleozol®, following Macherey Nagel’s protocol, and whole genomic DNA was isolated according to the phenol-chloroform purification and ethanol precipitation method of Sambrook, et al. [57]. Library construction (TruSeq RNA Library Preparation kit for RNA-seq and Swift Bioscience Accel-NGS Methyl-Seq DNA library Kit for WGB-seq) and high-throughput sequencing steps (Paired-End (PE) 2 × 150 bp, Illumina® HiSeq®) were performed by Genewiz (Leipzig, Germany). We performed all the bioinformatic preprocessing analyses using publicly available software implemented in the European UseGalaxy server [58].

#### Methylation calling

The WGB-seq raw reads were checked for quality using *FastQC*. Adapter and low-quality sequences were then trimmed using *Trim Galore!* Version 0.4.3.1. As specified by the Accel-NGS Methyl-seq Kit manual, additional 15 bp and 5 bp were also trimmed at the 5’ and 3’ extremity, respectively, to remove the tail added during library preparation and thus avoiding non-quality-related bias. Mapping was performed on the same references genomes as in RNA-seq analyses, using *Bismark Mapper* version 0.22.1 [59]. Only uniquely mapping reads were retained and PCR duplicates were removed using *Bismark Deduplicate* tool. We then extracted the methylation status from the resulting alignment files using *MethylDackel* (Galaxy Version 0.3.0.1), where only cytosines covered by a minimum of 10 reads in each library were considered, and with the option of excluding likely variant sites (i.e. minimum depth for variant avoidance of 10×, and maximum tolerated variant fraction of 0.95). In a previous study, we showed that cytosines at CpG context displayed the highest methylation levels (and variation thereof) in *Dunaliella salina* [23 and Table S2]. Furthermore, CpG-methylations have been suggested to play a role in gene regulations [60-62], and proposed as a molecular mechanisms underlying phenotypic plasticity [15, 17]. We thus investigated the genomic DNA methylation patterns in response to salinity across the different evolutionary treatments for cytosines at the CpG context. The high bisulfite conversion rate (> 99%) was assessed by Genewiz, by spiking in unmethylated lambda DNA in three randomly chosen libraries.

#### Gene expression analyses

The RNA-seq raw reads were checked for quality using *FastQC* version 0.72 [63] and subjected to adapter trimming and quality filtering using *Trim Galore!* version 0.4.3.1 [64]. Additional 12 bp and 3 bp were also removed at the 5’ and 3’ extremity, respectively, to avoid bias not directly related to adapter sequences or basecall quality according to *FastQC* outputs, and only reads with a minimum length of 50 bp were retained. We used the reference nuclear (Dunsal1 v. 2, GenBank accession: GCA_002284615.2), chloroplastic (GenBank accession: GQ250046) and mitochondrial (GenBank accession: GQ250045) genomes of *D. salina* strain CCAP 19/18 (closely related to CCAP 19/15 used here) for trimmed reads alignment, using *HISAT2* version 2.1.0 [65] with default parameters for PE reads and spliced alignment option. We finally quantified the number of reads per transcript with *FeatureCounts* version 2.0.1 [66] using the alignment files from *HISAT2* and the *de novo* transcript annotation produced for this species from Leung, et al. [23].

### 4.3. Statistical analyses

#### Differential DNA methylation and gene expression

We used Bioconductor’s *methylKit* package [67] to identify differentially methylated regions (DMRs), that is, non-overlapping 100bp windows with methylation levels that varied significantly among evolutionary treatments, or between salinities for each treatments. The significance of calculated methylation differences was determined using Fisher’s exact tests. We used the Benjamini-Hochberg (BH) adjustment of *P*-values (FDR < 0.05) and methylation difference cut-offs of 20%. Similarly, the differential gene expression analyses were performed with the Bioconductor’s package *DESeq2* version 1.30.1 [68]. We identified differentially expressed transcripts among the evolutionary treatments, the salinity, and their interaction, by building a general linear model as implemented in *DESeq2*. We used the Wald test when comparing two salinities, and transcripts with FDR < 0.05 (*P*-values after BH adjustment) and |log_2_FC| > 1 were considered as differentially expressed. Significance of evolutionary treatments (predictability *ρ*^*2*^) and *ρ*^*2*^ × salinity interaction term was assessed using likelihood ratio tests (LRT) comparing models with and without the interaction term [68].

We then used the Gene Ontology (GO) assignments from Leung, et al. [23] to classify the functions of *D. salina* transcripts and to functionally annotate the identified DE transcripts. Enriched GO terms of the DE transcripts were identified using *topGO* R package [69] classic algorithm and based on *p*-value generated using Fisher’s exact method, for the three GO categories, i.e. Molecular Function, Cellular Component and Biological Process. GO terms were then sorted by decreasing order of evidence within each category, based on the GO enrichment test *P*-value after Benjamini-Hochberg (BH) adjustment, and showed the most probable gene function candidates with a threshold of FDR ≤ 0.1.

#### Evolution of the degree plasticity at multiple levels

Since phenotypic plasticity is an adaptation to fluctuating and predictable environment [3-7], we wished to quantify to what extent environmental predictability during populations evolution contributed to key mechanisms underlying phenotypic plasticity. We first applied a redundancy analyses [RDA, 70], computed with the function *rda()* from the *vegan* R package [71], to quantify the proportion of the total epigenetic and gene expression variation that are significantly explained by the evolutionary treatments (predictability *ρ*^*2*^) and assay environment. We performed a RDA using the table of DNA methylation levels of 100bp windows regions, or the table of *rlog* transformed transcript count, as response variables, and the evolutionary treatments (predictability *ρ*^*2*^), salinity, and the *ρ*^*2*^ × salinity interaction, as explanatory variables, with population identity as covariates to account for paired samples between salinities. We treated the target *ρ*^*2*^ (low, intermediate or high) and salinity (low or high) as categorical variables. We quantified the contribution to the total genetic variation using the adjusted *R*^*2*^, and tested its significance by ANOVA-like permutation tests using 999 randomizations of the data [72]. For both RNA-seq and WGB-seq data, we presented the results of the RDA ordinations as a biplot, to visualize the variation among samples along its major axes. We performed all multivariate statistical analyses with the *vegan* R package [71].

To investigate how the overall degree of plasticity at multiple levels evolved in our experiment, we followed the protocol described in Leung, et al. [5], which we here applied to DNA methylation level, gene expression and cells morphological variation. (Note that the morphological data was here replicated, rather than just reproducing values from Leung, et al. [5]). First, we assessed the degree of plasticity of each experimental population by computing the Euclidean distance between the multivariate means from the CpG-methylation level table, the *rlog* transformed transcript count table, or the raw cells morphological data measured, at low ([NaCl] = 0.8 M) versus high ([NaCl] = 4.0 M) salinity. The realized autocorrelation of a given time series can vary to some extent from its long-term stationary expectation, because of the randomness of the stochastic process in finite time. For a more quantitative relationship between plasticity and environmental predictability, we thus used the realized (rather than the target) *ρ*^*2*^ as index of environmental predictability [as also done in 5]. For each time series, we thus calculated the realized environmental autocorrelation *ρ* as the correlation between salinities at two subsequent transfers. We then tested whether the degree of plasticity evolved according to environmental predictability, by regressing the Euclidian distance of plastic changes against the realized *ρ*^*2*^. The linear regression *t-Test* was applied to determine whether the slope of the regression line differs significantly from zero.

## Statement of authorship

L-MC, DG and CL designed the study. CL and DG performed the experiments and collected the data. CL analyzed all the data and prepared the figures and tables, with input from LQ for the epigenomic analysis. CL wrote the first drafts of the paper with L-MC. All authors reviewed and approved the final draft of the paper.

## Data and material availability statement

Raw sequence data (RNA-seq and WGB-seq) used in this study are deposited in the NCBI’s Sequence Read Archive (SRA) database under BioProject ID PRJNA736997. The specific samples used in this study are under the BioSample accessions listed in Tables S1 and S2, in supplementary materials.

## Acknowledgments

This work was supported by the European Research Council (Grant 678140-FluctEvol) to LMC, a Fonds de Recherche du Québec - Nature et Technologies (FRQNT) postdoctoral fellowship to CL, and a travel grant for collaboration provided by the GDR Plasticité Phénotypique (GDR 3715) from CNRS.

## Supplementary information

### Supplementary tables

**Table S1.** WGB-seq mapping statistics. For each sample, population ID, realized *ρ*^*2*^, and salinity are detailed in Table S2. We used a genome size of 350 M bp to estimate depth of coverage. Total C’s analysed were after reads deduplication and number of methylated C’s and percentage of methylation are given by cytosine context.

| BioSample<br>accessions | Total PE<br>reads | Estimated<br>depth of<br>coverage (×) | PE reads after<br>trimming (%) | Alignment<br>with a unique<br>best hit | Dupli-<br>cation rate | Total<br>C's<br>analysed | Methylated C's |  |  | % methylation |  |  |
| --- | --- | --- | --- | --- | --- | --- | --- | --- | --- | --- | --- | --- |
|  |  |  |  |  |  |  | CpG | CHG | CHH | CpG | CHG | CHH |
| SAMN19677484 | 53.59 M | 45.93 | 53.13 M (99.16%) | 20.13 M | 12.83% | 624.79 M | 34.90 M | 8.48 M | 5.00 M | 41.56% | 7.13% | 1.19% |
| SAMN19677485 | 47.54 M | 40.75 | 47.01 M (98.88%) | 17.34 M | 11.95% | 542.37 M | 31.66 M | 8.53 M | 4.47 M | 43.83% | 8.18% | 1.22% |
| SAMN19677486 | 47.03 M | 40.31 | 46.61 M (99.10%) | 17.84 M | 12.88% | 536.34 M | 29.00 M | 7.46 M | 4.31 M | 39.63% | 7.42% | 1.19% |
| SAMN19677488 | 50.90 M | 43.63 | 50.22 M (98.65%) | 19.33 M | 13.97% | 574.38 M | 32.50 M | 1.39 M | 4.43 M | 42.41% | 1.27% | 1.14% |
| SAMN19677489 | 51.80 M | 44.40 | 50.84 M (98.15%) | 19.33 M | 18.78% | 532.67 M | 29.98 M | 7.48 M | 4.54 M | 41.85% | 7.41% | 1.26% |
| SAMN19677490 | 48.41 M | 41.50 | 47.22 M (97.53%) | 16.59 M | 13.55% | 496.96 M | 28.95 M | 7.39 M | 4.28 M | 44.14% | 7.67% | 1.28% |
| SAMN19677492 | 55.62 M | 47.67 | 55.32 M (99.46%) | 19.92 M | 11.10% | 693.64 M | 45.23 M | 8.25 M | 5.61 M | 51.33% | 5.90% | 1.20% |
| SAMN19677493 | 48.79 M | 41.82 | 48.52 M (99.46%) | 17.44 M | 10.26% | 620.39 M | 40.25 M | 7.17 M | 5.16 M | 51.86% | 5.69% | 1.24% |
| SAMN19677494 | 51.23 M | 43.91 | 50.85 M (99.26%) | 18.78 M | 12.09% | 600.03 M | 36.14 M | 7.62 M | 4.86 M | 45.86% | 6.53% | 1.20% |
| SAMN19677499 | 52.76 M | 45.23 | 52.14 M (98.82%) | 19.96 M | 14.77% | 609.52 M | 32.61 M | 8.19 M | 4.94 M | 39.55% | 7.14% | 1.20% |
| SAMN19677500 | 57.53 M | 49.31 | 56.99 M (99.07%) | 21.43 M | 14.48% | 655.66 M | 36.42 M | 9.30 M | 5.27 M | 41.52% | 7.43% | 1.19% |
| SAMN19677501 | 54.14 M | 46.40 | 53.55 M (98.91%) | 20.36 M | 14.09% | 616.86 M | 33.66 M | 8.05 M | 4.73 M | 40.50% | 6.87% | 1.13% |
| SAMN19677503 | 43.92 M | 37.65 | 43.28 M (98.55%) | 16.90 M | 15.30% | 483.33 M | 25.42 M | 1.12 M | 3.62 M | 38.71% | 1.24% | 1.10% |
| SAMN19677504 | 53.14 M | 45.54 | 52.62 M (99.02%) | 19.75 M | 14.53% | 597.58 M | 32.95 M | 7.50 M | 4.68 M | 41.33% | 6.51% | 1.16% |
| SAMN19677505 | 54.72 M | 46.91 | 54.04 M (98.76%) | 17.80 M | 14.67% | 526.53 M | 28.74 M | 7.74 M | 4.52 M | 40.29% | 7.70% | 1.27% |
| SAMN19677507 | 50.07 M | 42.92 | 48.91 M (97.68%) | 17.44 M | 15.49% | 481.82 M | 25.15 M | 6.62 M | 4.25 M | 38.24% | 7.33% | 1.31% |
| SAMN19677508 | 46.82 M | 40.13 | 44.91 M (95.92%) | 15.31 M | 16.26% | 423.57 M | 23.08 M | 5.86 M | 3.77 M | 40.31% | 7.20% | 1.32% |
| SAMN19677509 | 47.91 M | 41.06 | 46.93 M (97.97%) | 16.95 M | 13.91% | 503.94 M | 27.33 M | 6.53 M | 4.32 M | 40.54% | 6.80% | 1.27% |
| **Populations founded from a single cell:** |  |  |  |  |  |  |  |  |  |  |  |  |
| SAMN19677483 | 55.29 M | 47.39 | 54.46 M (98.50%) | 20.10 M | 13.98% | 596.41 M | 32.82 M | 9.31 M | 4.74 M | 40.50% | 8.26% | 1.18% |
| SAMN19677487 | 47.81 M | 40.98 | 47.24 M (98.79%) | 17.97 M | 13.01% | 551.24 M | 32.13 M | 1.31 M | 4.32 M | 44.04% | 1.24% | 1.16% |
| SAMN19677491 | 54.56 M | 46.76 | 53.85 M (98.70%) | 20.75 M | 13.83% | 626.74 M | 34.69 M | 8.91 M | 5.59 M | 40.86% | 7.53% | 1.32% |
| SAMN19677498 | 52.72 M | 45.19 | 52.43 M (99.45%) | 19.37 M | 11.37% | 652.53 M | 39.01 M | 9.22 M | 5.33 M | 45.95% | 7.14% | 1.21% |
| SAMN19677502 | 44.98 M | 38.56 | 43.82 M (97.42%) | 17.55 M | 17.89% | 467.65 M | 22.14 M | 1.02 M | 3.50 M | 33.85% | 1.21% | 1.10% |
| SAMN19677506 | 53.95 M | 46.24 | 53.10 M (98.43%) | 20.23 M | 16.56% | 563.74 M | 28.33 M | 7.90 M | 4.70 M | 36.56% | 7.49% | 1.23% |

**Table S2.** RNA-seq mapping statistics.

| BioSample<br>accessions | Population<br>ID | Realized<br>$\rho^2$ | Salinity | Total PE<br>reads | PE reads after<br>trimming (%) | Overall<br>alignment<br>rate |
| --- | --- | --- | --- | --- | --- | --- |
| SAMN19677484 | C0021 | 0.001 | 0.8 M | 22.10 M | 21.97 M (99.43%) | 76.01% |
| SAMN19677485 | C00111 | 0.000 | 0.8 M | 21.12 M | 20.99 M (99.43%) | 75.76% |
| SAMN19677486 | C0041 | 0.002 | 0.8 M | 23.18 M | 23.10 M (99.66%) | 75.84% |
| SAMN19677488 | C09101 | 0.791 | 0.8 M | 24.11 M | 24.00 M (99.54%) | 75.43% |
| SAMN19677489 | C0951 | 0.720 | 0.8 M | 20.57 M | 20.48 M (99.58%) | 75.41% |
| SAMN19677490 | C0981 | 0.616 | 0.8 M | 23.64 M | 23.52 M (99.49%) | 75.32% |
| SAMN19677492 | CM05101 | 0.272 | 0.8 M | 22.43 M | 22.30 M (99.42%) | 75.51% |
| SAMN19677493 | CM05151 | 0.267 | 0.8 M | 21.83 M | 21.68 M (99.31%) | 75.21% |
| SAMN19677494 | CM0571 | 0.223 | 0.8 M | 29.63 M | 29.49 M (99.54%) | 75.83% |
| SAMN19677499 | C0021 | 0.001 | 4.0 M | 24.01 M | 23.89 M (99.53%) | 76.50% |
| SAMN19677500 | C00111 | 0.000 | 4.0 M | 22.44 M | 22.35 M (99.59%) | 76.07% |
| SAMN19677501 | C0041 | 0.002 | 4.0 M | 24.24 M | 24.14 M (99.57%) | 76.37% |
| SAMN19677503 | C09101 | 0.791 | 4.0 M | 27.00 M | 26.90 M (99.60%) | 76.39% |
| SAMN19677504 | C0951 | 0.720 | 4.0 M | 21.35 M | 21.25 M (99.54%) | 76.58% |
| SAMN19677505 | C0981 | 0.616 | 4.0 M | 22.67 M | 22.57 M (99.60%) | 76.35% |
| SAMN19677507 | CM05101 | 0.272 | 4.0 M | 27.80 M | 27.68 M (99.55%) | 76.86% |
| SAMN19677508 | CM05151 | 0.267 | 4.0 M | 26.56 M | 26.44 M (99.57%) | 76.85% |
| SAMN19677509 | CM0571 | 0.223 | 4.0 M | 26.74 M | 26.64 M (99.65%) | 77.14% |
| **<u>Populations founded from a single cell:</u>** |  |  |  |  |  |  |
| SAMN19677483 | C0021_B1 | 0.001 | 0.8 M | 21.11 M | 20.99 M (99.47%) | 75.52% |
| SAMN19677487 | C09101_B1 | 0.791 | 0.8 M | 18.73 M | 18.64 M (99.51%) | 75.52% |
| SAMN19677491 | CM0510_D4 | 0.272 | 0.8 M | 24.08 M | 23.96 M (99.53%) | 75.25% |
| SAMN19677498 | C0021_B1 | 0.001 | 4.0 M | 22.52 M | 22.42 M (99.54%) | 76.69% |
| SAMN19677502 | C09101_B1 | 0.791 | 4.0 M | 22.42 M | 22.31 M (99.54%) | 77.04% |
| SAMN19677506 | CM0510_D4 | 0.272 | 4.0 M | 22.00 M | 21.90 M (99.56%) | 76.62% |

### Supplementary figures

**Fig. S1.**
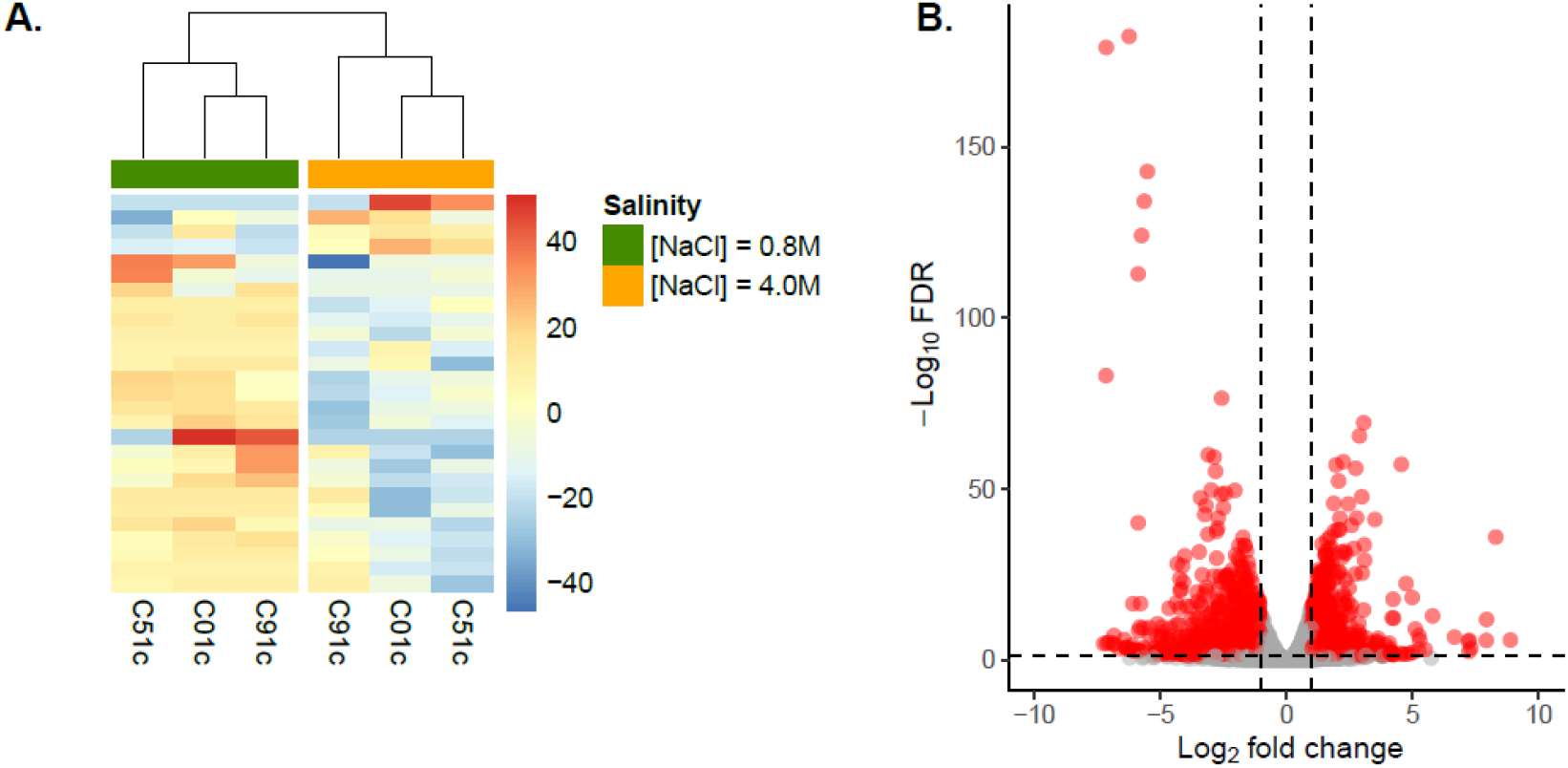
Salinity effect on DNA methylation and gene expression in isogenic populations. We founded three populations from single isolated cells from three evolved populations [following the protocol in 5]. As *D. salina* is haploid, a population founded from a single cell is expected to be isogenic. A. Heat-maps of WGB-Seq analysis for DMRs between salinities (n = 27). Each row represent a DMRs and column names are the population identity. Relative DNA methylation levels vary from blue (under-methylated) to red (over-methylated), as shown on the right-hand side of the heat-maps. Dendrograms on the top resulted from a hierarchical clustering analysis using the Euclidean distance of DNA methylation level among populations. B. Volcano plot illustrated significant (for FDR < 0.05 and |Log_2_FC| > 1) and non-significant differentially expressed transcripts between salinities as red and grey points, respectively. Salinity effect was assessed by comparing isogenic three populations (i.e. found from a single cell).

